# A Novel Cell Culture Scratch Assay Platform Generates Reproducible Gaps for Quantitative Cell Movement-Based Studies

**DOI:** 10.1101/2024.03.11.584450

**Authors:** Nicholas Wolpert, Lauren Gollahon

**Affiliations:** Department of Biological Sciences, Texas Tech University, 2500 Broadway, Lubbock, Texas 79409 US

## Abstract

Scratch assays are routinely performed for multiple research applications in cell biology (i.e., migration, metastasis, repair mechanisms, etc.). Conventional scratches are usually generated with pipette tips. User inconsistencies and varying pipette tips are major obstacles to reproducibility and quantitative power. Here, we present a novel, scratch/gap generating, cell culture plate-based, platform that addresses these issues, providing consistency and reproducibility across multiple users, significantly reducing variability, and increasing quantitative outcomes.

## Introduction

The rationale for developing this novel, scratch generating platform was the need to consistently reproduce gaps between users, into which, cell migration could be accurately quantitated for studies in metastatic breast cancer ^1^. Scratch assay or wound healing assay are different names for the same methodology ^2-4^. These assays serve a variety of purposes, (i.e., investigating cell migration behavior and speed, environmental effects, etc.). Basically, an area within the monolayer of cultured cells is removed and the remaining cell movements are observed. There are varying techniques by which a “scratch” or gap can be generated, scoring the surface of a culture dish with a pipette tip, cell scraper, toothpick, scalpel, or an expensive scratch assay machine ^5,6^. These manual (and frequently used) techniques vary significantly depending upon individual. Thus, reproducibility is greatly reduced due to inconsistencies in width and depth of the scratches. To gain quantitative power, specialized and expensive instrumentation, generally outside the operating budget for labs, is required. This instrumentation is also cost prohibitive to labs branching into studies of cancer cell metastasis, cell migration or repair ^7^. The focus of this project was the development of a device to address these inconsistencies, providing reproducible scratch widths and depths, across different users and well plate sizes (6, 12, and 24 wells). The goal was to decrease standard deviations while increasing quantitative and statistical power.

## Methods

### Development of the Device

The device invented was designed to create the same scratch each time and reduce the risk of error between users. Furthermore, it can be modified for petri dish use by using a different base. This scratch device is inexpensive and entirely customizable to the users’ preferences; including the ability to change the width of the tip to accommodate longer experiments or a larger wound, as well as the length of the tip to change the working distance, based on cell type. The default width for most 1 mL pipette tips is about ∼0.5mm. The device was designed to accommodate tips ranging in size from 500 microns up to 4 mm by using an interchangeable system that does not impact the procedure or configuration of the device. This was accomplished by employing fused deposition modeling, utilizing polypropylene as an autoclavable material to ensure sterility for aseptic conditions ^8,9^. Additionally, ethylene oxide sterilization can be used in the tip fabrication if repeated sterilization is necessary ^8^. Furthermore, the reproducibility of the device surface scoring among both researchers experienced in scratch assays as well as first time performers of the method was demonstrated. US Provisional Patent Application # 63471160 was filed June 5, 2023, under title: Wound Assay Device And Related Methods Of Use.

### Design Process and 3D Printing

The base of the device which goes over the plate (6, 12, or 24 wells) was originally designed to create flexible railing system with markers along the rail so that the user could pick where in the well to make the marks however making living hinges with the rigidity necessary to provide strength enough to lock the system in place and also be accurately placed proved to be difficult with the materials at hand. Instead, a rigid and fixed rail system which would allow the user to easily use the rail bar which goes across the well plate, or petri dish was used instead. The very first iteration of this design was for a 100mm petri dish and can be seen in Figure 1.

**Figure 1.**
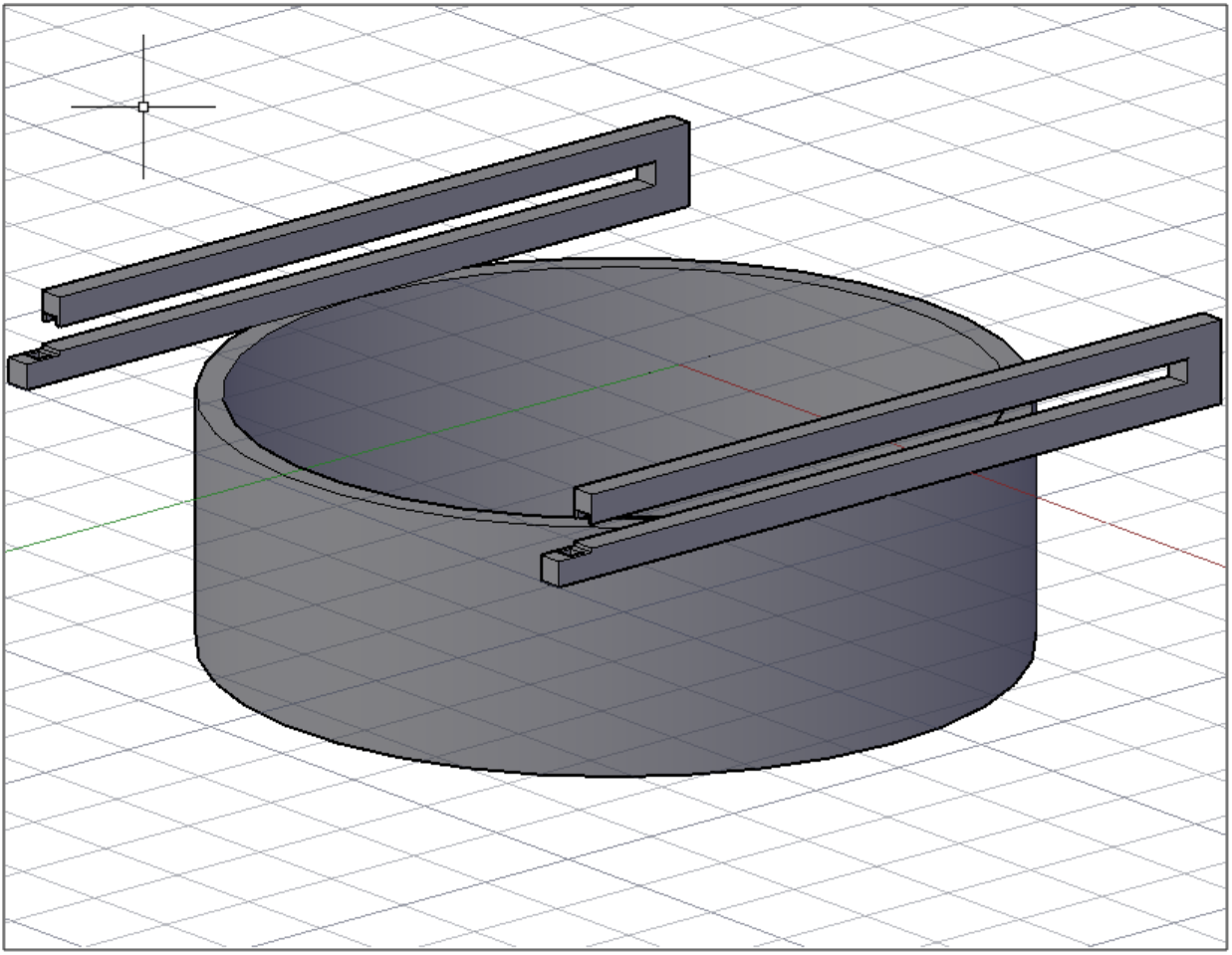
First cell scrap design with sliding rails for 100mm petri dish. The railing design for this too flimsy and caused too much sway in the scratching tip when in use.

Some of the iterations of the 6 well plate design can be seen in Figure 2. The railing system was designed to create a singular scratch on each well right in the center of the well for each well on the plate. The current platforms are made for a 6 well, 12 well, and 24 well plate. If needed before seeding the plate can be flipped over and the device can be used to trace a line for visibility on the reverse side. The plates were modeled in AutoCAD after measurements were taken by hand and the device base was designed around this modeling.

**Figure 2.**
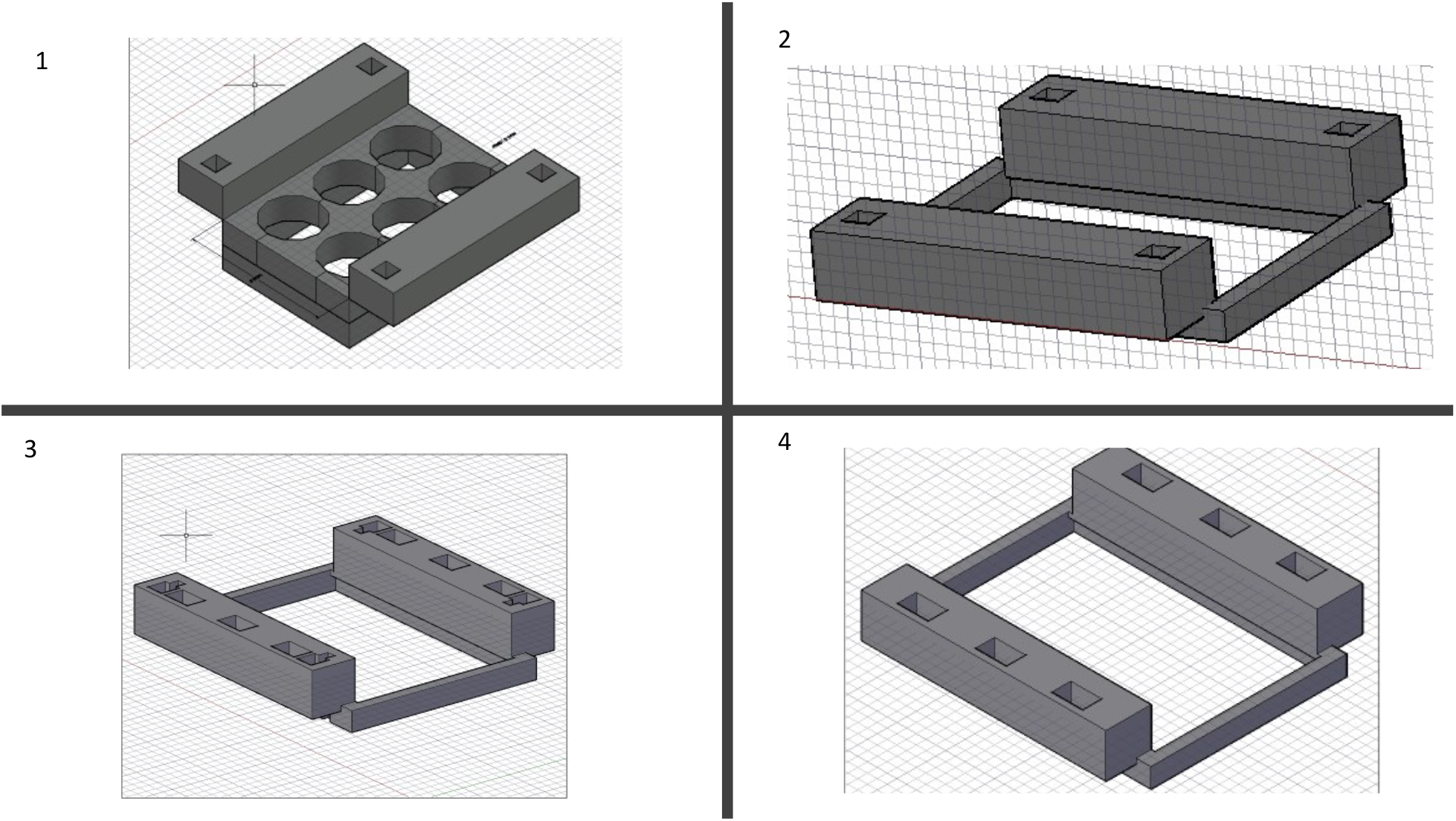
Design iterations in chronological order. Beginning with 1 which was intended to provide a larger shell for better accuracy for where the well is, 2 removed the plate guides making it more universal and easier to find the wells, 3 add divots for a proposed locking mechanism to hold down the railings and finally 4 is the current design for the 6 well plate where the holes provide guidance for the center of the wells of the plate.

The fabrication of the device was done using a DREMEL DigiLab 3D45-01 printer printing at the ultra-high resolution with an infill of 15% with support generated as necessary. Through the CURA software provided with the DREMEL printer.

The device was created to be user-friendly, while addressing potential inconsistencies that are naturally created between users i.e., pushing down to hard, different scratch lengths, jagged edging due to shaking hands, or creating crooked, uneven lines across the surface. The design includes a rail system in which the rail is removable and interchangeable to each of the subsequent plate bases that were created using the 6, 12, and 24 well plate sizes. The rail design means that the center of the scratch will always be in the center of the well. The plate base snaps onto the well plate to create a snug fit with no room for movement.

The design process involved looking at the current methods for creating scratch assays for small to mid-sized labs or labs just testing out scratch assays for their current research projects. In this regard the tip was originally designed to mimic a pipette tip by measuring the opening at the end of a 1mm pipette tip so that the line achieved would be the same width with the key distinction of being a more rigid tip to reduce the risk of deforming the tip with pressure from the user creating wider and thinner portions of the line as its being used. Creating a more rigid design involved giving the tip a longer base but maintaining the same width resulting in tip width that could be scaled up or down if needed to increase the or decrease the time necessary to close the wound.

The fabrication of the device tip was done using a DREMEL 3D printer printing at the ultra-high resolution with an infill of 100% with no supports and any holes generated from the .stl file being sliced was auto filled using the CURA software for model slicing.

The tips were designed using an oval shape which tapers into square base leaving behind a curved geometry that allows the scratching tip to remove the cells in a single pass. Previous design iterations of the tip can be seen in Figure 3, with the final design in box 2 (right panel).

**Figure 3.**
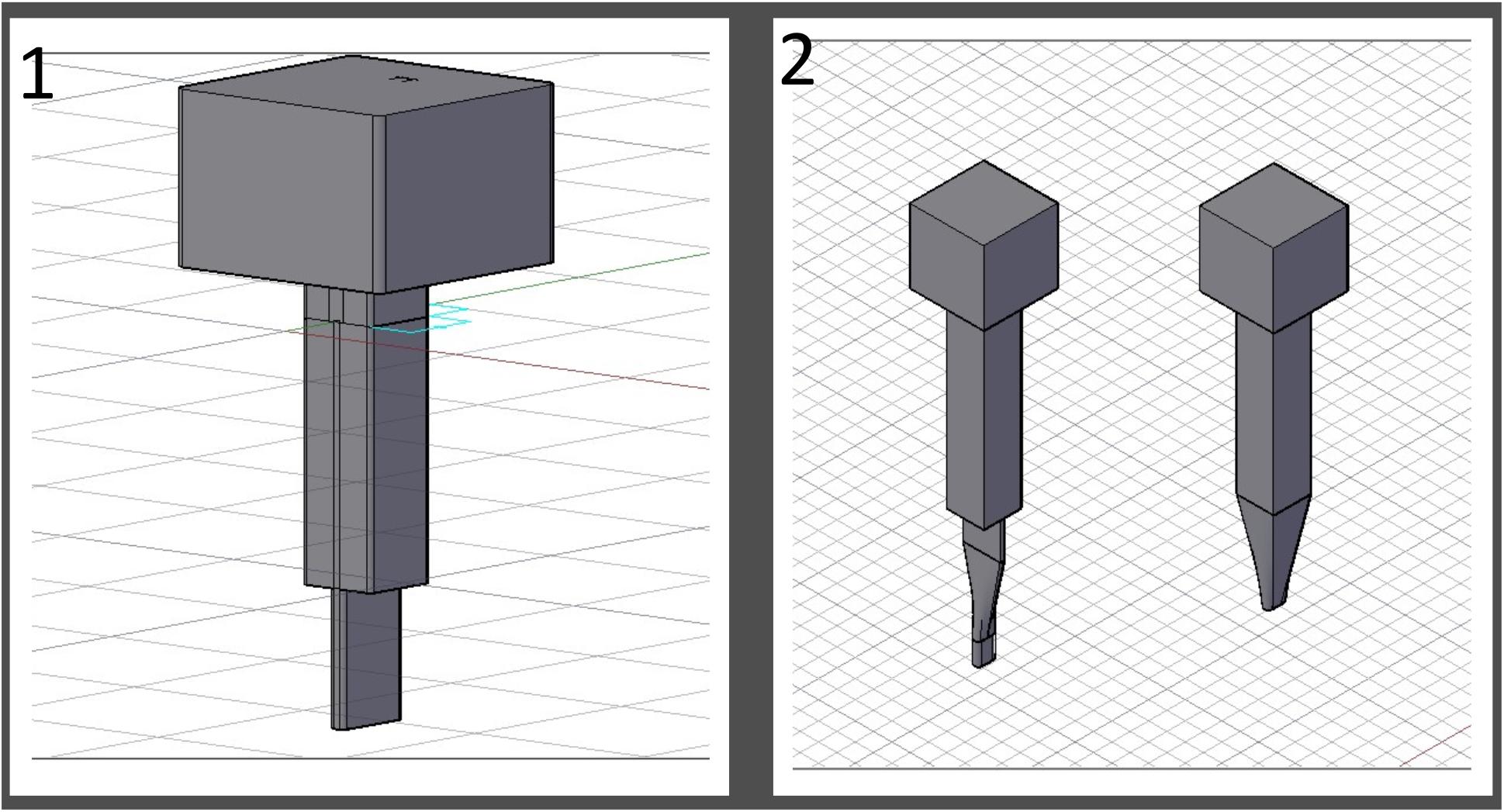
Previous design iterations of the tip. Starting with 1 which was the first design and was unstable and hard to print given the long unsteady protrusion, next was the tip on the left in box 2 which had similar geometry issues and finally the tip on the right in box 2 was the most stable and provided the perfect intersection of oval base with enough stability to print with relative ease.

This design shape makes it less likely for double lines to form if the user goes over the same area twice in a back-and-forth motion. It also ensures a straighter boundary will be formed from the score, making image analysis more consistent and reproducible.

### Testing the Device

The following procedure was used for growing the cells to test the device. For purposes of proof-of-principle, MCF7 triple positive MCF7 breast cancer cells (ATCC HTB-22) were used as the test cell line because our laboratory research focus is primarily breast cancer. However, this device can be used for any adherent cell type. To characterize attributes of the device, MCF7 cells were cultured on Thermanox™ coverslips (FisherScientific 174985) in Dulbecco’s modified Eagle medium (DMEM), supplemented with 10% fetal bovine serum and 100 µg/ml penicillin-streptomycin solution, under humid conditions at 37°C and 5% CO_2_, to a confluency of ∼ 90%. Thermanox™ coverslips were used in place of glass to facilitate cutting samples in preparation for scanning electron microscope (SEM) imaging. Treated glass coverslips may also be used as well for scoring directly in the well. Scoring substrate is based on imaging system working distances as there might be slight differences between the substrate removed from a glass coverslip versus from a the bottom of a well or the substrate found on the Thermanox™ catalog number 174985 coverslip which are coated with cell culture substrate. The Thermanox™ coverslips are also not recommended for fluorescent work as autofluorescence can be seen in the 380nm to 540nm range. But they work great for scanning electron microscopy processing.

### Similarities and differences between pipette and device tip

The device tip was designed to be the same width as the 1mL pipettes which we have in our lab which are the Fisherbrand SureOne ^tm^ non-filter graduated pipette tips with a filling capacity from 100 - 1250µL catalog number 02-707-400. The measurement was taken using calipers to find the width of the round tip which was 0.5mm in diameter. From this the device tip was then designed to create a scratch within this parameter. Whereas the pipette tip has a purely circular area of contact the device tip has oval area of contact in order to increase the area of contact with the surface in an effort to decrease the amount of pressure that would be pushed into the plate with a width 0.5mm and a length of 2mm the device tip with the same amount of force applied as the pipette tip would experience half as much force across the given area than the pipette tip at least with force acting in the downward direction or z plane which is demonstrated in Equation 1 and can be seen in Figure 4.

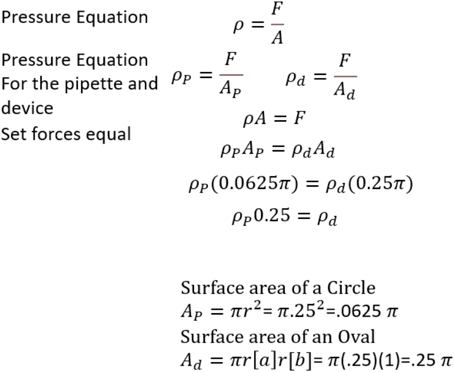

*Equation 1. Transformation of pressure equation to demonstrate z-plane force distribution*.

**Figure 4.**
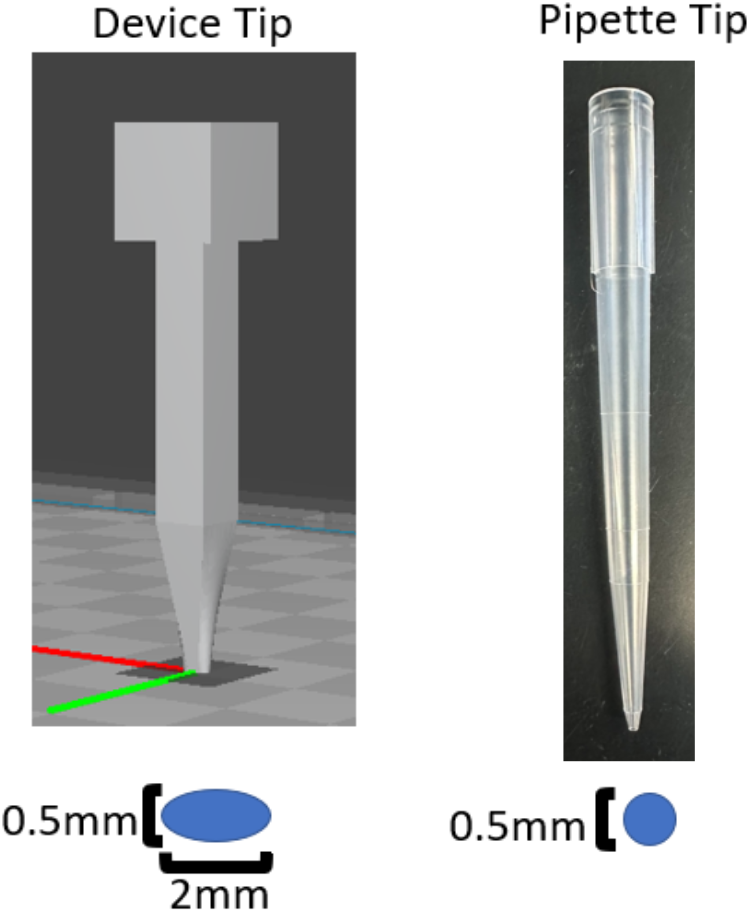
Comparison between the device tip and the pipette tip with the cross sections of the end of the tip beneath them.

### SEM Imaging Preparation

A Zeiss Crossbeam 540 scanning electron microscope was used at the same magnification and as similar as possible working distance in order to ensure the width sizes could be accurately compared across all samples along with these settings the electron high tension (EHT) for the microscope was kept as consistent as possible while providing a clear image. SEM imaging was done as opposed to fluorescent to be able to see a larger field of view of the scratch while being able to obtain higher resolution images for data collection.

SEM Preparation occurred immediately after the scratch assay was performed, the coverslips were fixed using a modified methodology previously described by Fischer in preparation for SEM imaging ^10^. The protocol adapted to work with these adherent cells is as follows:

1. Thermanox™ coverslips, were placed coated side up in a 6-well plate, for cell culture, and wells filled with appropriate cell culture medium.
2. MCF7 cells were seeded according to well-plate size (i.e., 6, 12, 24) and grown to 90% confluency.
3. Once confluency was reached the scratch was applied either with a 1 mL sterile pipette tip or using the device and accompanying tip.
4. After scratch application, the medium was aspirated, and the cells washed with PBS.
5. Once PBS was removed, the following steps were performed under a fume hood. There, 0.5ml of 2.5% glutaraldehyde in 0.1M sodium cacodylate buffer (CAC) was added to each well slowly to avoid causing the coverslip to float.
6. Cells were fixed at 4°C overnight in the 6-well plate wrapped in parafilm.
7. The samples were then washed with 0.1M CAC, 3 times for 2 min each time.
8. The post fixation was performed in a solution of 1% OsO_4_ in dH_2_O for 45min.
9. Then the samples were rinsed for 2 min with 0.1M CAC followed by 2 more washes, for 2 min each, with dH_2_O.
10. These washes were followed by an ethanol (ETOH) dehydration series.
  a. 25% ETOH, once for 5 min
  b. 50% ETOH, once for 5 min
  c. 75% ETOH, once for 5 min (stored overnight at 4°C)
  d. 95% ETOH once for 1 min
  e. 100% ETOH 3 for 5 min
11. A 1:1 mix of (Hexamethyldisilazane (HMDS): ETOH) was used for 10min.
12. Two 100% HMDS bathes at room temperature for 10 min
13. Excess liquid was removed by capillary flow by placing a paper towel to the edge of the coverslip. Samples were then dried for at least 4 hours.

The final step was sputter coating with the Q150T ES plus EMS Quorum. The Thermanox coverslips were coated with a 4nm layer of iridium and imaged with the Zeiss Crossbeam 540 SEM.

## Results

Consistency and reproducibility was verified by SEM, comparing the device to manual methods of scratch formation. This scratch device is inexpensive and entirely customizable to the users’ preferences; including changing tip width and length to accommodate experimental times, conditions and working distance, based on research questions. The default width for most 1 mL pipette tip brands is ∼0.5 mm. By using an interchangeable system design, the device accommodated tips ranging in size from 0.5 mm to 4 mm. This was accomplished by employing fused deposition modeling, utilizing polypropylene (autoclavable), ensuring sterility for aseptic conditions ^8,9^.

For device validation, MCF7 breast cancer cells (ATCC HTB-22) were used due to our laboratory research focus in breast cancer. However, this device can be used for any adherent cell type. To determine reproducibility among and between users, MCF7 cells were cultured on Thermanox™ coverslips (FisherScientific #174985) as described in the on-line Methodology section, to facilitate cutting samples in preparation for scanning electron microscope (SEM) imaging analysis (shown in Figure 5). The device tip was 3D printed to generate a 0.5 mm width. SEM imaging was performed to concurrently examine a larger field of view while acquiring higher resolution images for data collection and analysis.

**Figure 5.**
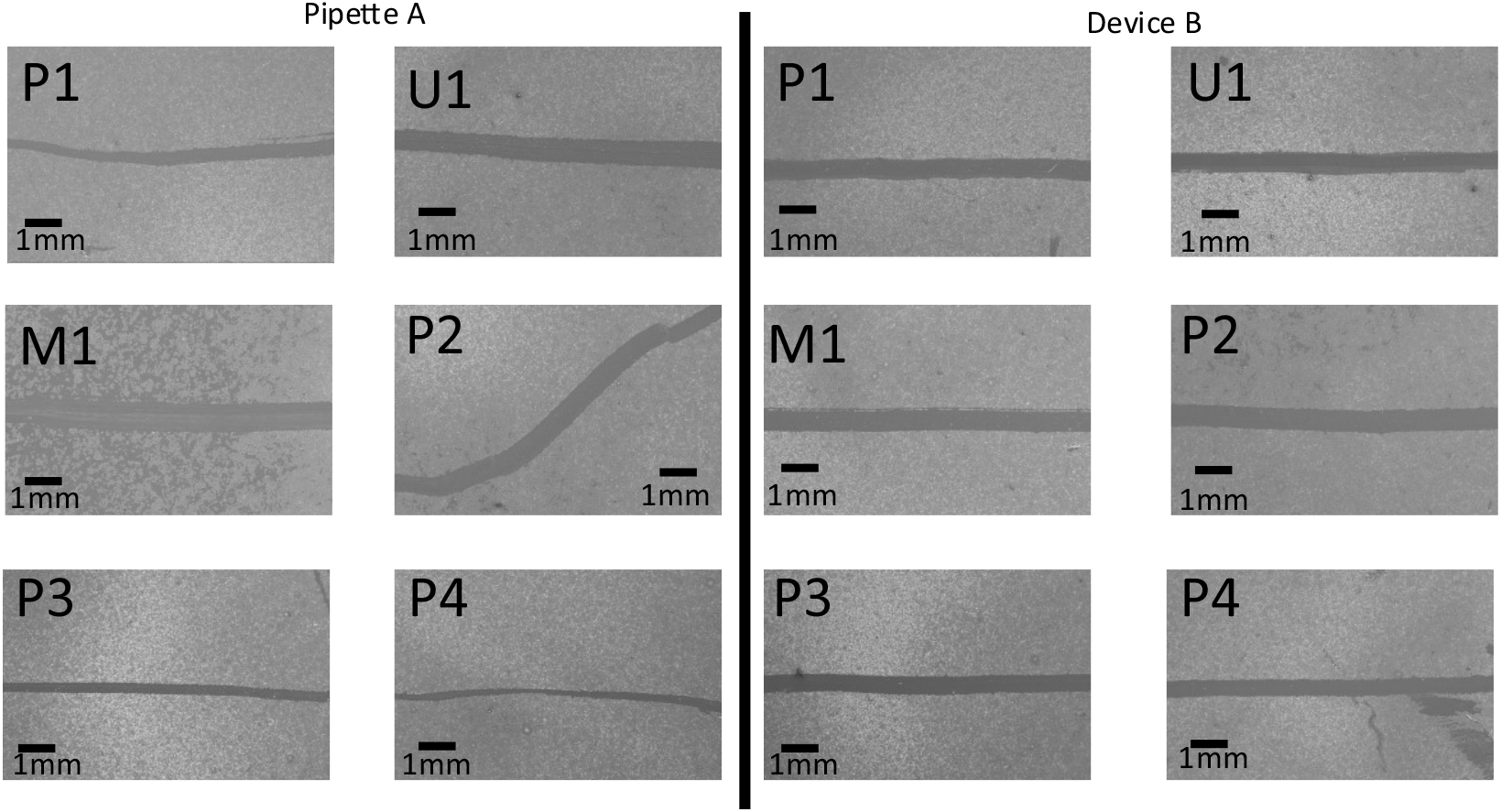
Pipette scratch (A) and device scratch (B) imaged with a Zeiss Crossbeam 540 scanning electron microscope. The same individual generated scratches as labeled. EHT = 15.00kV, 12X Magnification, working distance: 47.4mm.

At ∼80 – 90% confluency, cell cultures were scored by 6 laboratory personnel designated P1 – P3 (PhD students), M1 (Master’s student), and U1 (undergraduate). SEM preparation was performed immediately after scoring the cell culture dish. Coverslip SEM preparation follows Fischer ^10^ with modifications for these adherent cells. Figure 5 illustrates the SEM results, post-fixation, for cell scratches created by the device and pipette tip per user. Scratch widths were measured at different distances along their lengths to determine consistency and reproducibility, analyzed and compared using FIJI ^11^. This involved taking the threshold value to create a binary image and creating preliminary differentiation of the background layers from the scratch for accurate measurements. The distances were standardized using the same pixels per millimeter ratio across all images. Next, the function *fill holes* was used to clean up the image and removed small aberrations to alleviate bias in the final analysis step, *Erode*, which was applied to remove small holes in the image, generating a cleaner environment for the final function. The last function and analysis were performed using the *local thickness* command along with its associated masked, calibrated, and silent settings. The distances were set by binning from 0.0 to 0.9 mm with 256 bins for the histogram. The width of the scratch created by the device averaged 0.571±0.0645 mm (n=364,921). In contrast, the pipette tip had an average width of 0.467±0.235 mm (n=301,756). Figure 6 depicts a histogram breakdown of the width differences for user P1.

**Figure 6.**
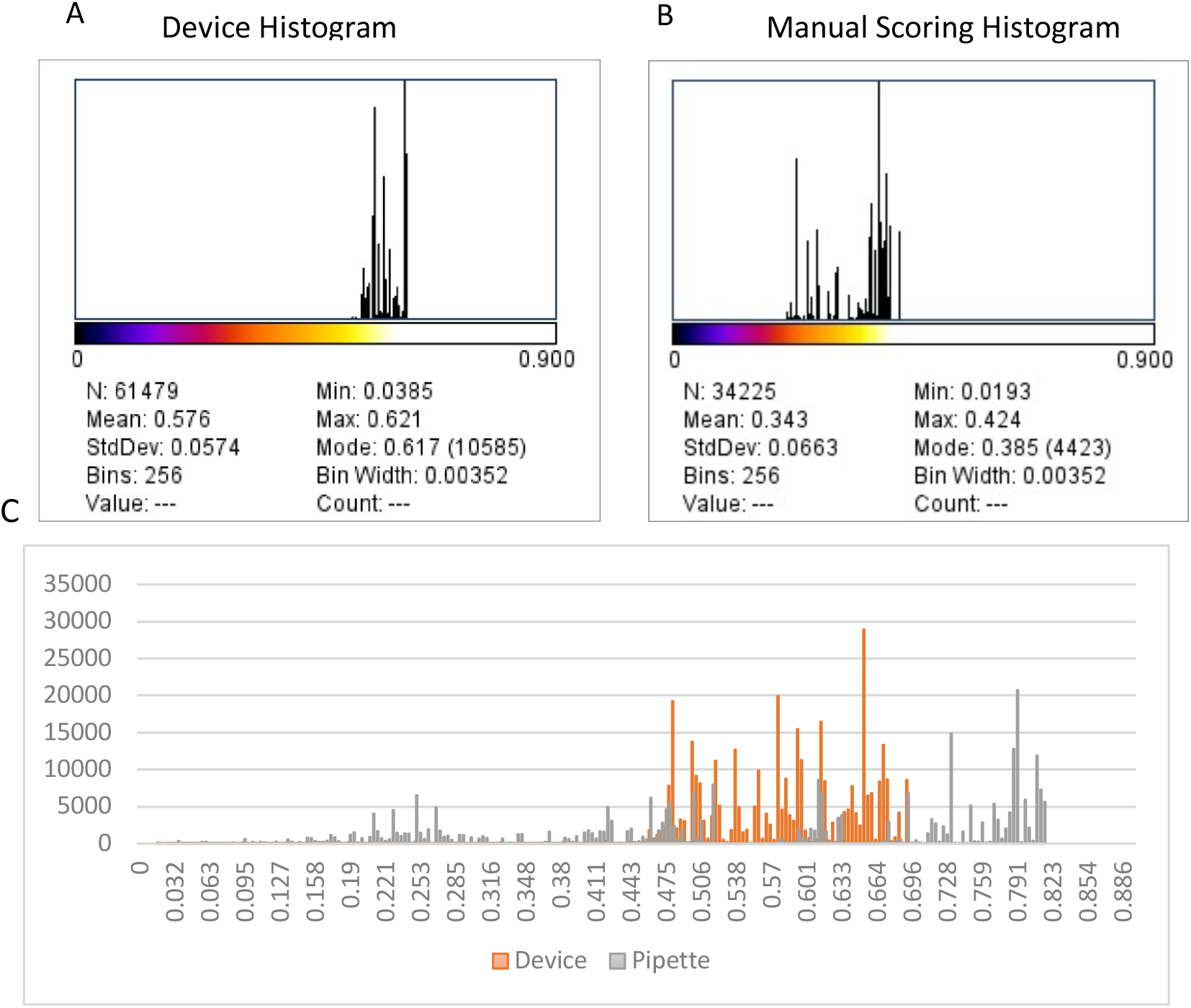
Comparison of device and user generated scratch width variability. A) Graphical representation of bins breakdown used for measuring the distance from 0 to 0.9mm. Mean value of 0.576mm for the radius = average width of the device achieved by user P1). B) The histogram from user P1 for manual scratch generation. C) Histogram of device and pipette data points in relation to their widths in mm for P1.

## Conclusions

Device design is modular, facilitating application to different cell culture platforms (6, 12, and 24 well plates). Tip width is customizable between 0.5 mm – 5 mm. Tip length can be modified based on working distance, cell and culture plate type, and components can be autoclaved. Device robustness allows consistent and reproducible operation between users and experimental replicates, along the whole length of the scratch, resulting in significantly reduced standard deviations and greater quantitative and statistical power. Overall, a highly significant difference at a confidence interval of 99.999 and a p value of p< 0.001, was observed between the manual (pipette) and device scratches. US Provisional Patent Application # 63471160.

## Author Contributions

Conceptualization, N.W. and L.G.; methodology, N.W.; formal analysis, N.W.; resources, L.G.; writing—original draft preparation, N.W.; writing—review and editing, L.G.; supervision, L.G. All authors have read and agreed to the published version of the manuscript.

## Acknowledgements

Gollahon Lab members, Dr. Karin Ardon-Dryer - real-time microscopy instrumentation, Dr. Bo Zhao - College of Arts Microscopy Center.

